# Reconstruction of in-vivo subthreshold activity of single neurons from large-scale spiking recordings

**DOI:** 10.1101/673046

**Authors:** Stylianos Papaioannou, André Marques Smith, David Eriksson

## Abstract

Current developments in the manufacturing of silicon probes allow recording of spikes from large populations of neurons from several brain structures in freely moving animals. It is still, however, technically challenging to record the membrane potential from awake behaving animals. Routine access to the subthreshold activity of neurons would be of great value in order to understand the role of, for example, neuronal integration, oscillations, and excitability. Here we have developed a framework for reconstructing the subthreshold activity of single neurons using the spiking activity from large neuronal populations. The reconstruction accuracy and reliability have been evaluated with ground truth data provided from simultaneous patch clamp membrane potential recordings in-vivo. Given the abundance of large-scale spike recordings in the contemporary systems neuroscience society, this approach provides a general access to the subthreshold activity and hence could shed light on the intricate mechanisms of the genesis of spiking activity.

## Introduction

In any species and in any neuronal circuit the membrane potential plays a key role in transforming and distributing neuronal signals. The membrane potential integrates incoming information, it influences the function of ion channels so they can modulate subsequent input, and, whether it crosses a threshold or not, it transmits this new information to other neurons (Stuart & Spruston 2015, Takahashi *et al.* 2016). In addition to its physiological importance the membrane potential is a continuous signal that is richer than a discrete signal such as a train of action potentials. Finally, the membrane potential can reveal fast changes in neuronal physiology which cannot be seen with a smoothed spike train. These can be the fast changes that occur during normal brain function or in response to an experimentally controlled perturbation.

Ideally one would like to measure the membrane potential of hundreds of neurons in multiple different areas and different layers in the brain, and in a freely moving animal of any kind. Such large-scale recordings of membrane potential are so far impossible. To measure the membrane potential there are electrophysiological methods using glass pipettes and imaging methods using fluorescent indicators (Wang *et al.* 2019). Electrical recordings using a glass pipette are regarded as “the gold standard” since they measure the membrane potential directly. Such recordings have been done in anaesthetized animals (Kodandaramaiah *et al.* 2012, Yu & Ferster 2010, Volgushev *et al.* 2006, Lampl *et al.* 1999), head fixed awake animals (Zhao *et al.* 2016, Gentet *et al.* 2010), and even in freely moving animals (Lee *et al.* 2006). To patch multiple cells in vivo is difficult and may in the future be facilitated by patch “robots” (Kodandaramaiah *et al.* 2018, Suk *et al.* 2017). Imaging methods might be more suitable for addressing the membrane potential of multiple cells simultaneously (Bando *et al.* 2019), but may still represent a smoothing of activity, owing to limitations on scanning and shuttering speed when compared to electrophysiological sampling rates. There is a strong effort to develop fluorescent indicator constructs that approximate the membrane potential as close as possible (Quicke *et al.* 2019, Piatkevich *et al.* 2018, Chamberland *et al.* 2017, Gong *et al.* 2015, Flytzanis *et al.* 2014, Zou *et al.* 2014, Hochbaum *et al.* 2014, St-Pierre *et al.* 2014, Gong *et al.* 2014, Jin *et al.* 2012), and an equally strong push in developing imaging tools to be able to image deep in the brain and even in the freely moving animal (Wang *et al.* 2018, Ghosh *et al.* 2011, Helmchen & Denk 2005).

To be able to approach large scale membrane potential recordings we propose to use large scale neuronal spiking activity to reconstruct the membrane potential of individual neurons. In fact, the physiology of the brain facilitates this thanks to the many nuances of redundancy, being it branching axons (Kalil & Dent 2014), gap junctions (Goodenough & Paul 2009), ephaptic input (Anastassiou *et al.* 2010), or recurrent/mono/poly-synaptic input. Because of this redundancy, a sub threshold event will occur in multiple neurons, some of which will generate a spike because of this event and others because of distinct input (**Figure 1**). In line with this redundancy it is possible to predict the spiking activity of one neuron given the spiking activity of other neurons (Truccolo *et al.* 2010).

**Figure 1.**
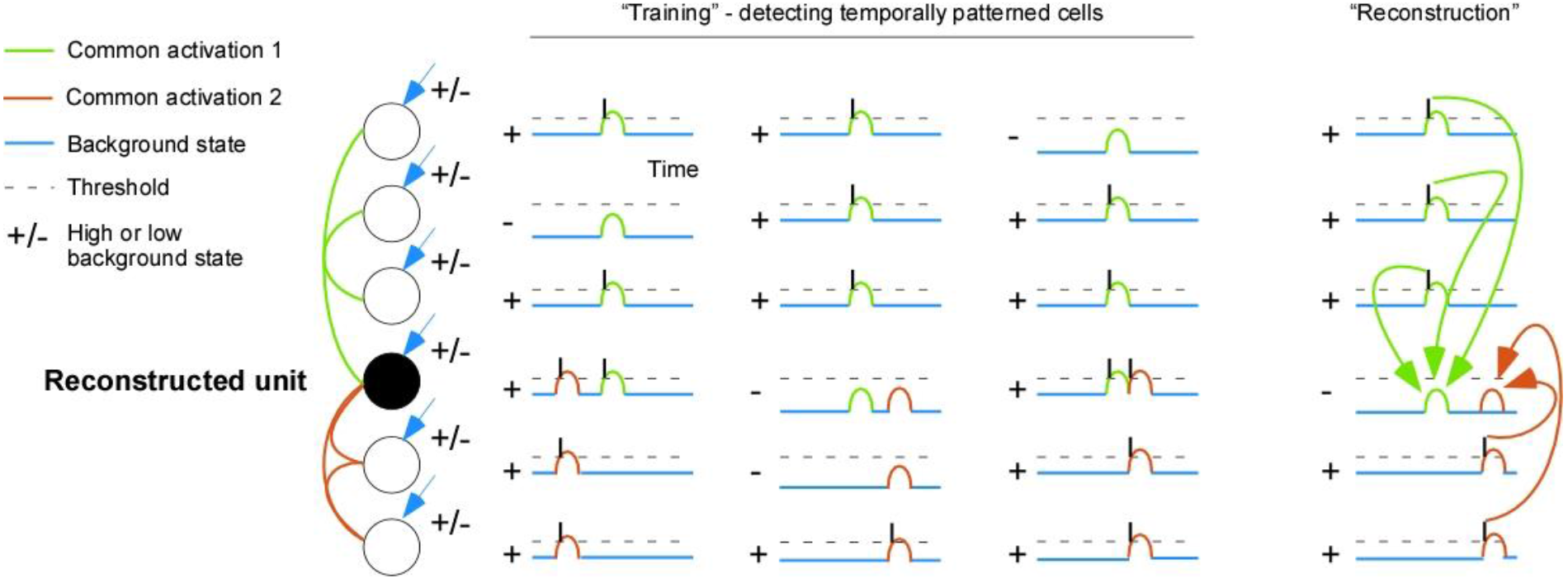
Using spatiotemporal spike patterns to reconstruct subthreshold activity. An artificial neuronal network is trained to find spatiotemporal patterns that are associated with the spikes of the reconstructed unit. During the reconstruction phase those associations are used to fill in subthreshold activity.

Here we test a framework for reconstructing subthreshold activity for a specific unit from a set of spike times and their unit identity (**Figure 2A**). We have developed a cost function that only penalizes the reconstruction when it is above the threshold at a time point where there is no spike (**Figure 2B**). Therefore, anything can happen below the threshold without it being corrected for. The reconstruction is done by an artificial neuronal network that detects spatiotemporal patterns in the population spiking activity. The framework is available at https://github.com/David-Eriksson/SubLab. Using data from simultaneous Neuropixel and intracellular recordings we show that the algorithm can reliably reconstruct the subthreshold dynamics. Then we show that the reconstructions are most reliable for de-synchronized spiking data. Finally, we use the reconstruction to reproduce some differences in membrane potential dynamics for anaesthetized and awake animals.

**Figure 2.**
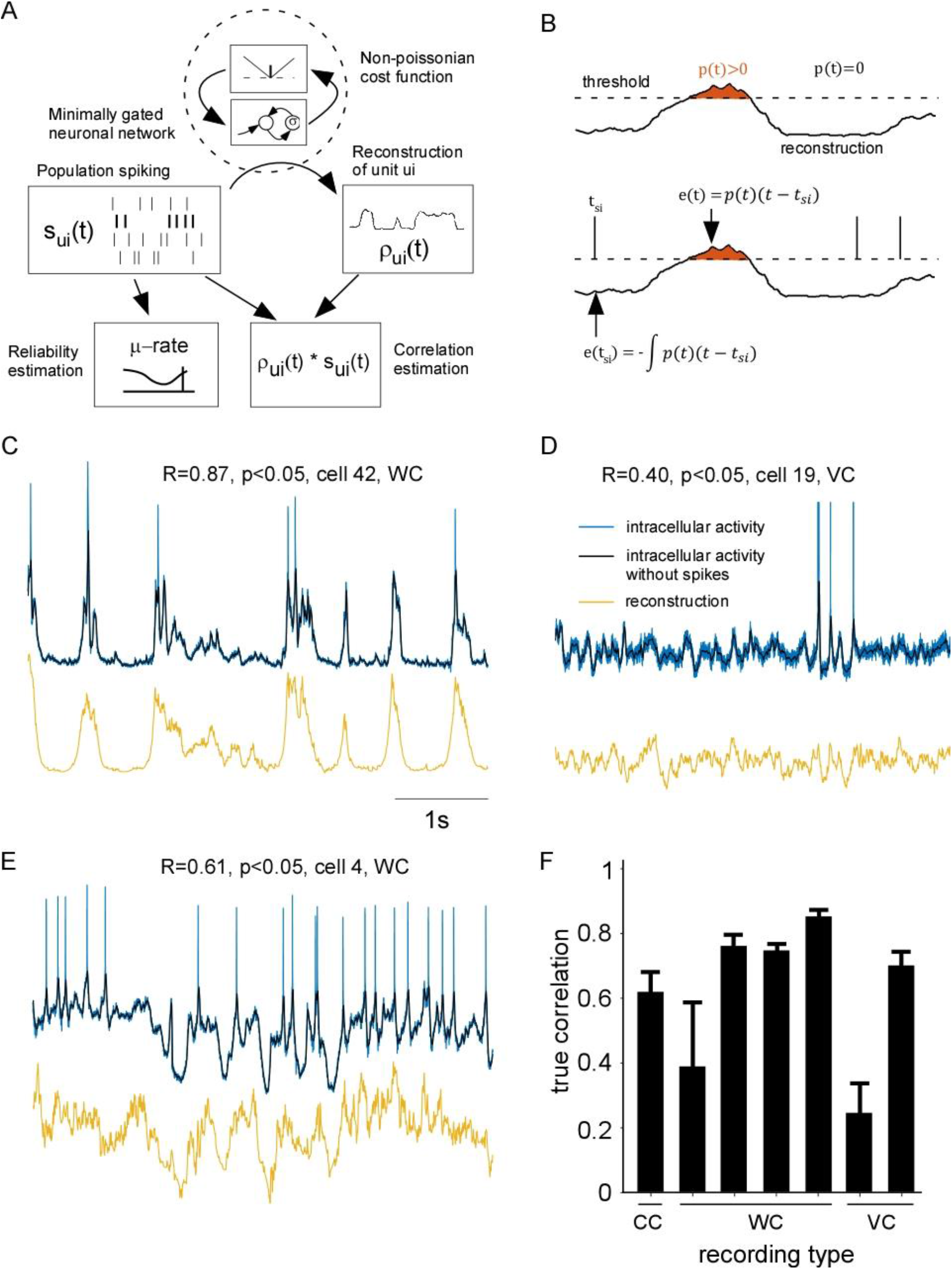
Verification of the reconstruction algorithm with data from membrane potential recordings in-vivo. **A**: Schematic of the subthreshold reconstruction framework (github: SubLab). **B:** Non-poissonian cost-function. The total number of spikes for the reconstructed neuron are distributed proportionally to the positive value of the reconstruction in order to create a spiking-probability, p(t) (top). This probability is then used to weight the spiking timing error between the true spikes and the spiking probability distribution of the reconstruction (bottom). The spike timing error can then be converted to one error per time bin that can be back-propagated in order to refine the reconstruction. **C:** Reconstruction (yellow) of the subthreshold dynamics of a cell and the actual whole cell recording (blue) from the same cell. The correlation between the two traces is noted on the top. **D**: The same as in **C**but for cell 19. The trace is inverted because the cell was recorded with voltage clamp. **E**: The same as in **C** but for cell 4. **F:** Distribution of the correlations between reconstructed and actual recordings across all 7 cells (CC=current clamp, WC=whole cell current clamp, and VC=voltage clamp). Each cell was reconstructed at 10 (except for the current clamped cell which only had 7 because of a shorter recording) different test intervals (as opposed to the remaining time points that served as training). This gave 10 (or 7) values for each cell for which the standard deviation was calculated.

## Results

### The algorithm reliably and consistently reconstructs the subthreshold activity of neurons from extracellular recordings

Here we apply the algorithm for reconstruction of membrane potential on a dataset of extracellular activity recorded with the Neuropixels probe (Jun *et al.* 2017) from the somatosensory cortex of the rat that was acquired in the Kampff lab (Marques-Smith *et al.* 2018a, Marques-Smith *et al.* 2018b). Depending on the session, between 50 and 200 units were identified using the Kilo-sort spike sorting algorithm (Pachitariu *et al.* 2016). An additional (target) unit was extracted from the simultaneous intracellular recording. Only intracellular recordings with a certain signal-to-noise ratio were included in the analysis (see Methods). The subthreshold activity of the target unit was reconstructed using the algorithm and was compared against the ground truth, the simultaneous current clamp or voltage clamp. Since the algorithm reconstructs subthreshold activity without knowledge of the spiking activity in the target neuron it is fair to estimate the correlation between reconstructed activity and the intracellular activity using all time points (including those where there are spikes). The algorithm can reconstruct the membrane potential of a whole cell recording during pronounced down/up-states with a correlation of 0.87 (**Figure 2C**). For a more desynchronized intracellular activity in voltage clamp the algorithm reconstructs peaks and troughs in the absence of spiking activity with a correlation of 0.4 (**Figure 2D**). For a third cell the more complex down/up state dynamics in a whole cell recording could be reconstructed with a correlation of 0.61 (**Figure 2E**). Overall the subthreshold dynamics could be reconstructed with an average correlation of 0.62±0.22 (**Figure 2F**).

Since a large contribution to the subthreshold activity originates from the spiking of neighboring neurons we investigated the correlation between average instantaneous firing rate across all units and subthreshold activity. Indeed, for a session with clear up/down states the instantaneous firing rate was correlated with the subthreshold activity (R=0.70) (**Figure S1A**). For this case the reconstruction increased the correlation by 0.2 (R=0.90). The largest percentual changes could be seen for desynchronized brain activity for which the reconstruction had 140% larger correlation than the mean instantaneous firing rate (**Figure S1B**). This accuracy gain is because the reconstruction is unit specific and as such the algorithm can filter out input units that are not relevant for the reconstruction of one specific unit. Overall, all cells except one could be better reconstructed using the algorithm than using the average firing rate (**Figure S1C**).

### Estimating reconstruction accuracy without membrane potential measurements

For experiments in which there is no access to the membrane potential one needs to approximate the reconstruction reliability in terms of the spiking activity exclusively. In this section we show that the “true” correlation between reconstruction and measured membrane potential can be approximated with a “spike only” derived correlation index. This index resembles the correlation between reconstruction and the corresponding spike times (see Methods). We also show how the discrepancy between those two correlations, i.e. correlation error, can be empirically estimated without the true correlation. In essence, data with repeating spatiotemporal spiking patterns will decrease the true correlation and increase the correlation error. To illustrate this we used a leaky integrate and fire simulation (see Methods). For a non-repeating spiking pattern the true correlations resembled those of the in-vivo data (**Figure 3A** and **B**), and the correlation index resembled the true correlation (R = 0.59, p<0.05, n=41) with a low correlation error (0.13±0.12, mean±std, n=35) (**Figure 3C**). On the other hand, when the spiking pattern is repeated every 4 seconds (i.e. single condition case) the true correlation decreased and the correlation index increased (**Figure 3D**). The true correlation decreases because there is not enough data (because of repeated data) for the algorithm to estimate the relation between subthreshold activity and spiking activity. Moreover, the correlation index increases because it is easier to learn to reconstruct repeating spikes than stochastic spikes. In general, the correspondence between the correlation index and true correlation increases with number of “conditions” (**Figure 3E**). By adding independent noise to each simulated neuron in the single condition the true correlation increases and the correlation error decreases (**Figure 3F and S2**). Thus, a large correlation error can be “saved” with added noise (**Figure 3G**).

**Figure 3.**
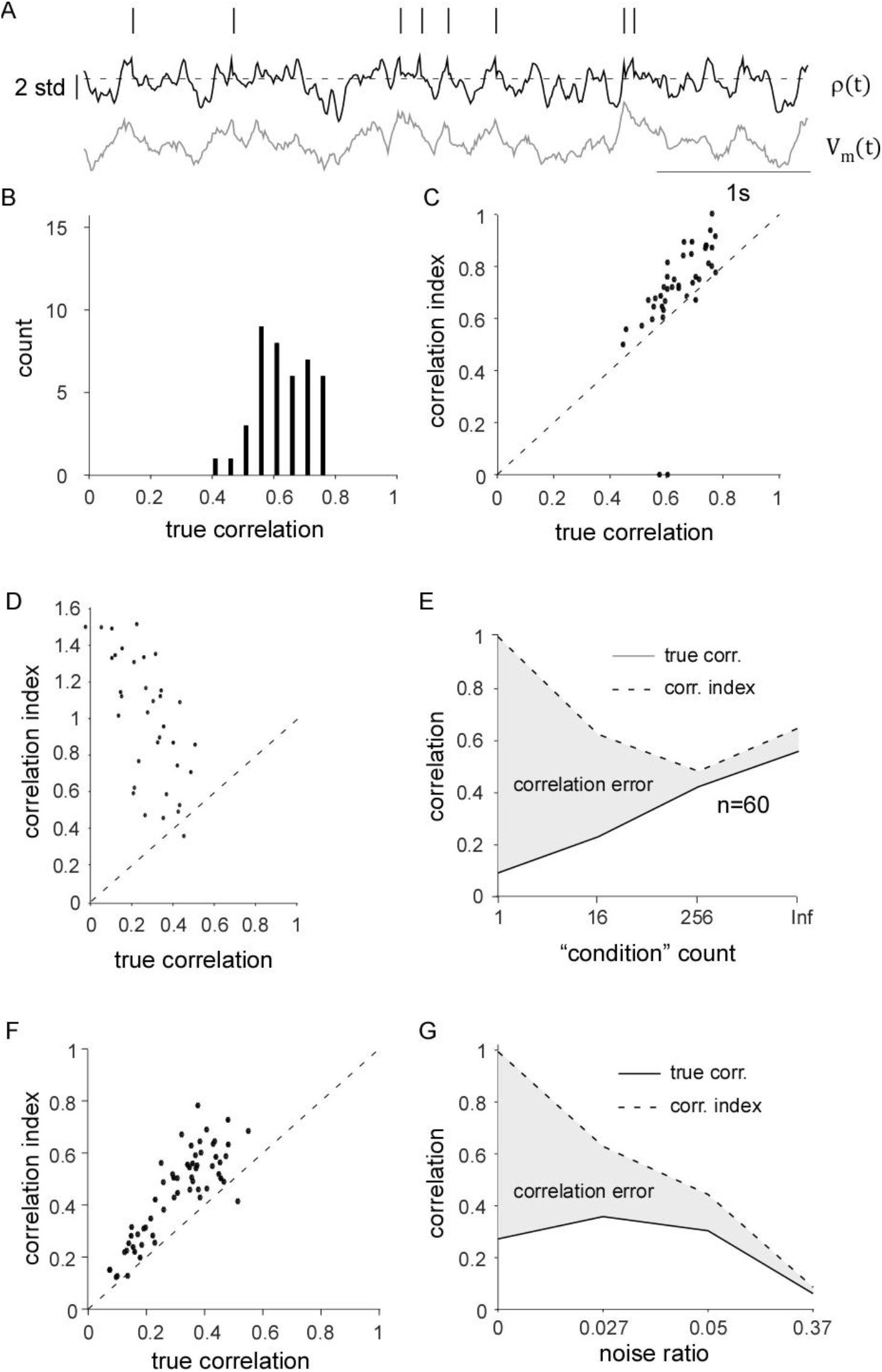
Simulated spiking repetitiveness decreases reconstruction reliability. **A:** Reconstruction, ρ(t), for an example trace (black) and the measured membrane potential V_m_ (gray) **B**: Distribution of the true correlations between ρ(t) and V_m_ for all simulated cells. **C**: Relation between correlation index which we call correlation index and true correlation for all cells for “infinite” number of conditions (low repetitiveness) **D**: Relation between correlation index and true correlation for the one condition case (high repetitiveness). **E**: The dependency on the number of conditions (degree of repetitiveness) and the estimated and true correlation. **F**: Relation between correlation index and true correlation for a noise ratio of 0.05. **G**: The dependency on the noise ratio and the correlation index and true correlation.

The repetitiveness can be described in general terms with the *μ*-*rate* (see Methods). It is maximal (average neuronal firing rate times number of neurons) for independent firing and it is zero when spikes cluster at a certain lag between the target neuron and input neurons (synchronization). Moreover, the *μ*-*rate* estimates the number of input spikes that is available for each time bin. For example, a μ-rate of 1000Hz corresponds to at least one input spike for each reconstructed millisecond.

The *μ*-*rate* increases as one increases the condition count which in turn decrease the correlation error from 0.3 to 0.07 (**Figure 4A**). Similarly, the *μ*-*rate* increases as one increase the noise ratio, which in turn decrease the correlation error by an order of magnitude (from 0.3 to 0.03) (**Figure 4B**). The *μ*-*rate* consistently separates reliable reconstructions from unreliable (**Figure 4C**). Accordingly, the correlation error decreases monotonically with an increasing *μ*-*rate* (**Figure 4D**). As a result, the correlation index estimates the true correlation within around ±0.1 if the *μ*-*rate* is more than 500 spikes per second. Since we have simultaneous intra-cellular measurements for the *in-vivo* data, we can now compare the correlation index with that of the true correlation (R=0.8, p<0.05, n=67). Like the leaky integrate and fire simulation the match between those two correlations depends on the *μ*-rate (**Figure 4E and F**). Here the correlation index estimates the true correlation within around ±0.1 if the *μ*-*rate* is more than 800 spikes per second. Thus the *μ*-*rate* is a general way to assess reconstruction reliability for both simulated data and *in-vivo* data for a large range of spiking statistics.

**Figure 4.**
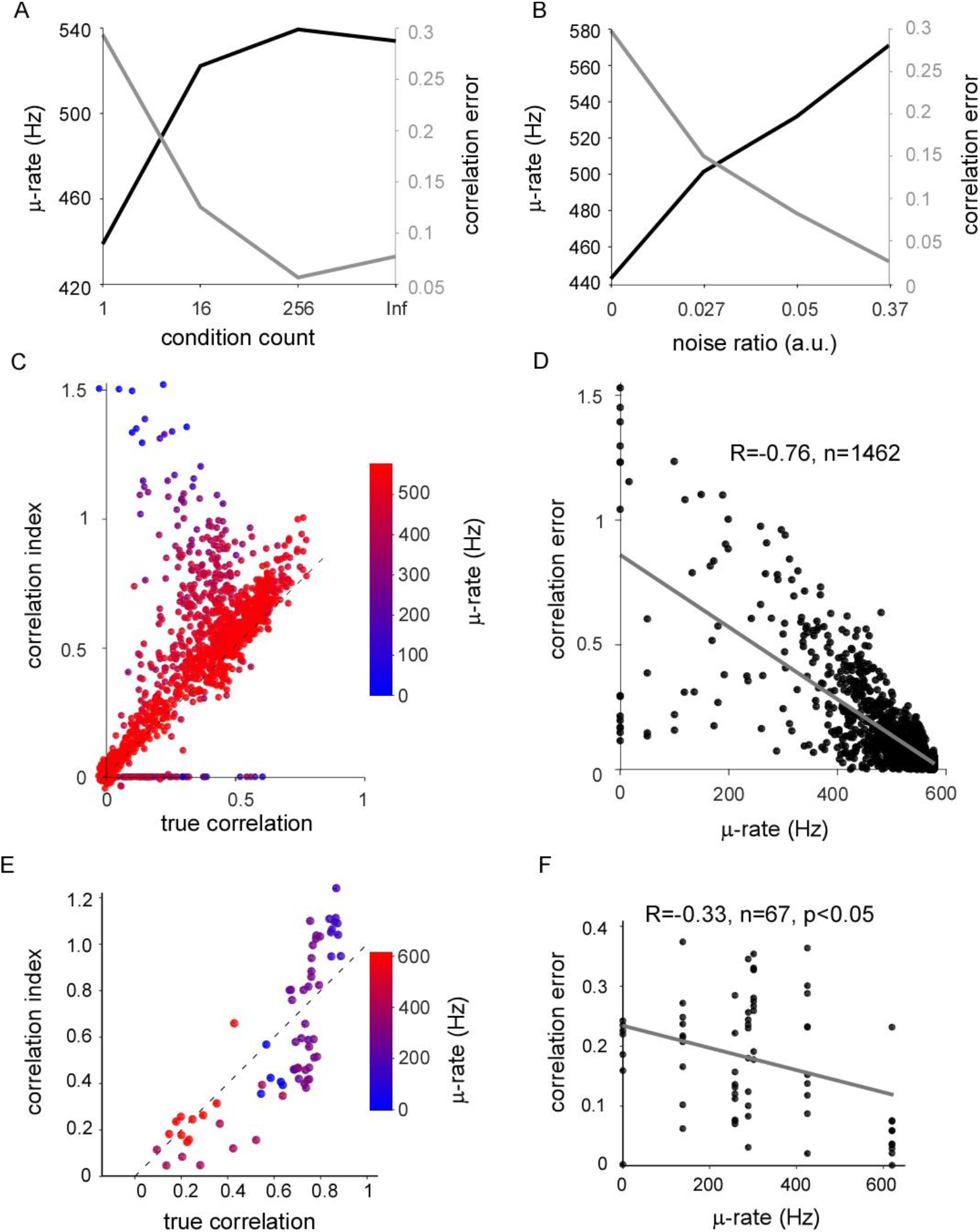
A single metric that predicts the reconstruction reliability. **A:** Relation between *μ*-rate, condition count, and correlation error. Note that for a specific condition count the *μ*-rate has been averaged across all noise ratios (see **Figure 3** and **S2**). **B:** Relation between μ-rate, noise ratio, and correlation error. Note that for a specific noise ratio the μ-rate has been averaged across all condition counts (see **Figure 3** and **S2**). **C:** Relation between correlation index and true correlation for different degree for different μ-rates. The data is pooled for all combinations of conditions (1, 16, 256, and Inf) and all noise ratios (0, 0.027, 0.05, and 0.37). **D:** Same as in C but as the relation between μ-rate and correlation error. **E:** Relation between correlation index and true correlation for different μ-rates for *in-vivo* data. **F:** Same as in **E** but as the relation between μ-rate and correlation error.

### Correlation index scales with unit count

In the future more neurons can be recorded at any given time. Therefore, we run the reconstruction algorithm with different number of neurons (**Figure 5A**). The data is from the motor cortex during spontaneous behavior of the awake mouse recorded by Dr. Nick Steinmetz at the UCL in the lab of Matteo Carandini and Kenneth Harris. We have used both single and multi-units for the reconstruction ending up with 471 units. On average the correlation scaled linearly with the logarithm of the unit count (**Figure 5B**). For the last step from 234 to 470 units the increase in correlation depended on the number of spikes, suggesting that the network is overfitted with fewer spikes (**Figure 5C**). Thus, although we choose this data set because it has the longest recording time it still does not seem to be enough data (spikes) to fit all the parameters for 471 units.

**Figure 5.**
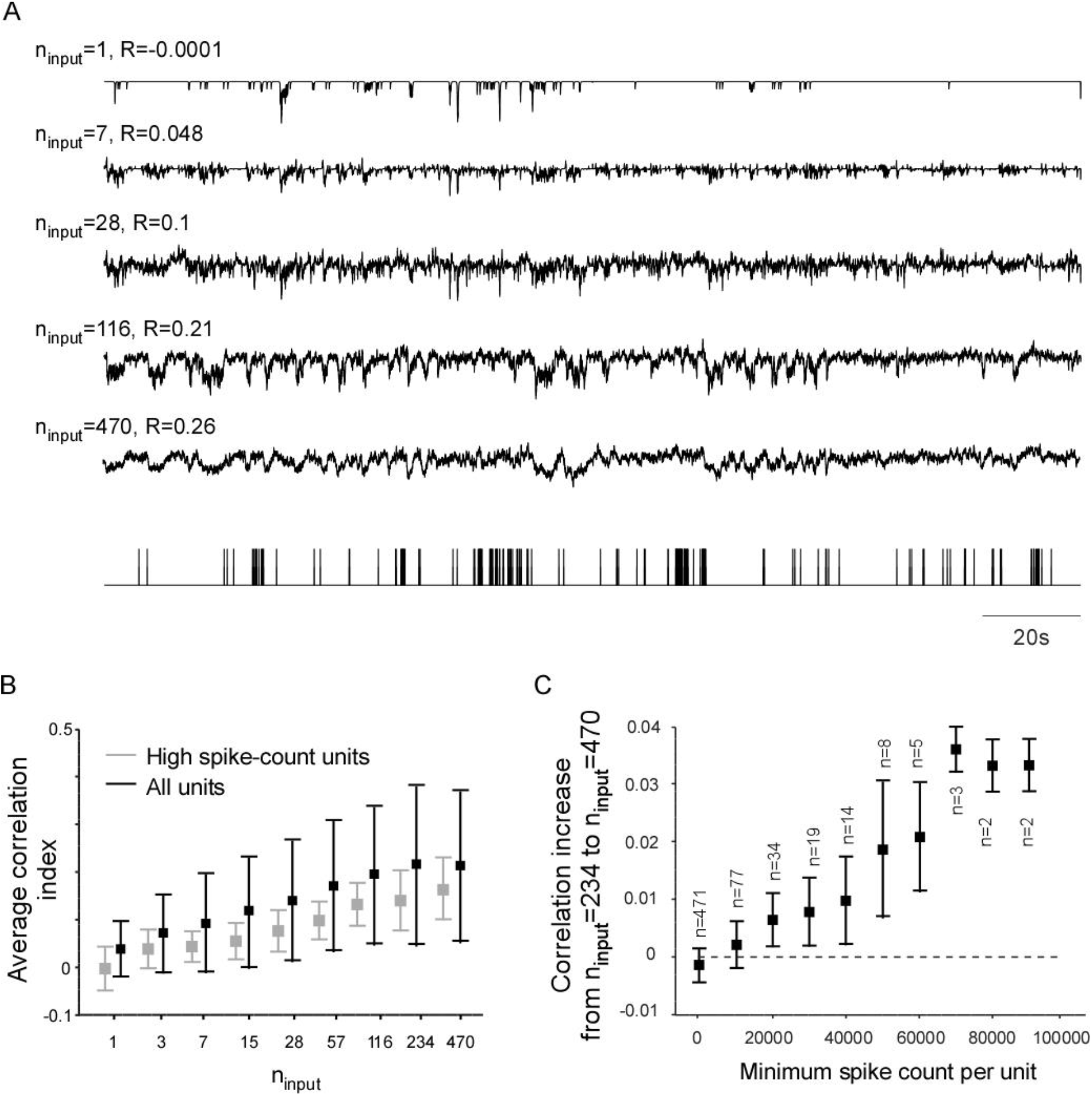
Correlation index scales with the number of units used as input **A:** Reconstruction of example trace for different number of units (five first rows). The target spikes were the same in all cases (bottom). **B:** The average correlation index for all units (black) and for high spike count units (gray) for different unit counts (error bars denote standard deviation). **C:** The increase in correlation from 234 to 470 units is dependent on the spike count of the target unit (error bars denote standard error).

### Opposing latency patterns across cortical layers in the awake and anesthetized animal

Here we have reconstructed the membrane potential of neurons from both anesthetized and awake recordings and studied their subthreshold laminar dynamics. We have used the Nick Steinmetz data set for the awake activity and the Kampff-lab data set for the anesthetized activity. To study the latency differences between cortical layers, the subthreshold activity was reconstructed using 1 ms resolution (**Figure 6A** and **B**). To study the latency across layers we detected all low to high transitions in the data set and averaged those (**Figure 6C** and **D**). The latency of a given unit was the time point when the activity crossed the 50% of the peak activity. In general, the latency decreased for deeper layers (−24±7ms/mm, M±SE) for the anesthetized case and increased (37±7ms/mm, M±SE) for the awake case (**Figure 6E**). This was also true for the 10% threshold, but not for the 100% threshold (**Figure S3A and B**). For the awake data there was not a significant correlation between the cortical depth and the 100% latency indicating a later synchronization when the activity becomes suprathreshold.

**Figure 6.**
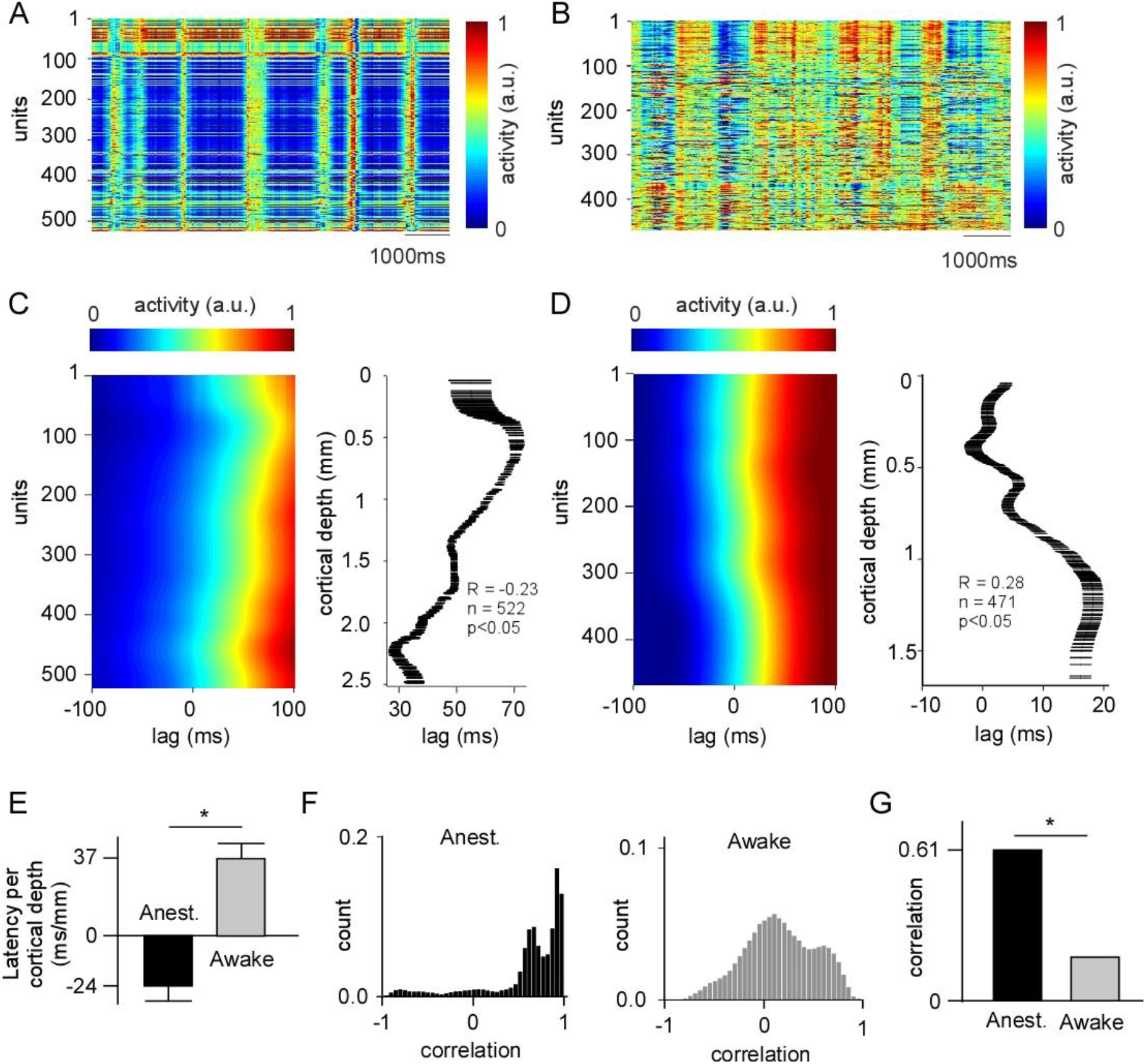
Comparison of reconstructed activity in the anesthetized and awake animal. **A:** Reconstruction of all 522 units from the anesthetized rat sensory motor-cortex, session 42 in the Kampff data base. **B:** Same as in A but for the 471 units from the awake recordings from mouse sensory-motor-cortex from the data base of Steinmetz. **C:** Rise triggered average of reconstructed activity (left). The lag is relative to the average (across units) threshold crossing time point. The activity was smoothed across neighboring units (see methods). Latency of the half-maxima for each unit (right). Bars indicate the standard deviation of the bootstrapped latencies. Note that the correlation statistics is for the latency as a function of cortical depth (not vice versa). **D:** Same as **C** but in the awake animal. **E:** Summary of the latency as a function of the cortical depth (mean±standard error). **F:** Distribution of cross-correlation values for all unit-pairs in the anesthetized animal (left), and for the awake animal (right). **G:** Average cross correlation in anesthetized and awake animal (the standard error is smaller than the printing resolution).

The correlation structure between neurons has been shown to depend on the state of the animal. Overall, the correlation between units was larger in the anesthetized state than in the awake state (**Figure 6F** and **G**). For both types of states, the correlation decreased towards infragranular layers (**Figure S3C** and **D**). This trend could also be seen in terms of the dimensionality of the population activity in each layer and by means of the visualized population trajectories (**Figure S3E** and **F**). Together this suggests that the latency of the subthreshold activity is different for the awake and the anesthetized state, but that the relative complexity between the layers remains the same for the two states.

## Discussion

Large scale spiking recordings is a universal technique that has been applied in all species ranging from *C-Elegans* to humans, anesthetized to freely moving animals. Here we present an algorithm that uses large scale spiking data to reconstruct subthreshold dynamics of individual neurons. We have verified the algorithm directly using in-vivo intracellular data. Moreover, analyzing the reconstructed subthreshold activity from awake and anesthetized recordings, we have further confirmed the validity of the reconstructed data by means of verifying already known opposing subthreshold dynamics across the cortical laminae for two discrete states, i.e. awake vs anesthetized.

The reconstruction algorithm implicitly relies on correlations between neurons to extract a continuous subthreshold activity from discrete spikes. Previously, correlations have been used to reduce dimensionality in order to estimate a smoothed firing rate using poissonian or gaussian statistics (Pandarinath *et al.* 2018, Cunningham & Yu 2014, Yu *et al.* 2009, Stopfer *et al.* 2003, Wu *et al.* 2017). Such an approach is, however, suboptimal for reconstructing subthreshold activity since the non-thresholded poissonian or gaussian statistics will cause the activity to vanish in the absence of spikes. Subthreshold activity may, however, be arbitrary large as long it remains below the threshold. Therefore, we have devised a thresholded non-poissoninan spike timing error function which is not correcting subthreshold activity in the absence of spikes. Due to the nature of dimension reduction techniques they typically need one or more parameters to set the number of dimensions and to control how the dimension reduction should be done for the data at hand. The reconstruction algorithm, on the other hand, is “plug-n-play” in the sense that there are no explicit parameters (it was not tuned to the particular data-sets used in this study). Furthermore, in contrast to dimension reduction techniques the reconstruction algorithm allows each neuron to have a unique nonlinear mapping to the other neurons. Thus, the algorithm could be defined as a dimension conserving technique for subthreshold reconstructions.

As a result of the dimension conservation the algorithm can only be applied if there are enough neurons and/or spikes for making the reconstruction (see **Figure 5C**). Although, here we have used data from the neuropixels probe to record an adequate number of neurons, more neurons could potentially be sampled for a longer time using thin-film electrodes (Chung *et al.* 2019) or with calcium imaging (Pachitariu *et al.* 2018). In the case of a brain area with small volume and few neurons it may become necessary to put additional electrodes in functionally related areas. Note that the algorithm is insensitive to whether the input area is sending axons to, or receiving axons from, the target area. Since the algorithm is acausal it can tune the reconstruction for either case or for both. Overall although the algorithm is dependent on a large-scale-recordings we expect the algorithm to be less restrictive as to where those recordings should be done in relation to the target neurons.

In order to reconstruct the subthreshold membrane potential, we rely on the naturally occurring asynchronicity of large-scale neuronal activity. This temporal jitter is captured by the μ-rate which is high when spikes are not occurring at a certain lag between two neurons. A large μ-rate indicates that there are enough “independent” data samples for the algorithm to find the relation between subthreshold and spiking events. The μ-rate is maximal when the neurons are firing independently of each other in which case the μ-rate becomes equal to the average neuronal firing rate times the number of simultaneously recorded neurons. A large μ-rate renders the reconstruction more accurate both for ground truth in-vivo data and for simulation data. For the in-vivo data, however, the correlation-index underestimates the true correlation. This could suggest that the spikes are not purely driven by the membrane potential of the neuron. There could for example be a noise component that is independent of the membrane potential, and independent between the neurons. Such a noise signal could be averaged out when taking multiple neurons into account as is done for the reconstruction. Indeed, for an independent noise process there is also an underestimation of the true correlation (**Figure S2**). All in all, both simulations and in-vivo recordings suggest that, for an accurate and reliable reconstruction, the spikes should not form repeating spatiotemporal patterns.

The non-repetitiveness in turn suggests that the reconstruction algorithm works best for non-overtrained animals or well-trained animals performing many different conditions. Importantly, this does not mean that the experiment becomes less interpretable. On the contrary, first, the functional connectivity between two signals, being it sensory-neuronal, neuronal-neuronal, and neuronal-behavioral is more reliably estimated if one of the signals (or both) is experimentally randomized (Eriksson 2016, David *et al.* 2004). This is because randomization typically cancels out long term correlations that would otherwise bias the estimate. Second, the brain will not “adapt” to a stereotypical behavior by changing the locus of representation to a another brain area (Kawai *et al.* 2015). Nevertheless, the algorithm has been able to reconstruct both synchronized and desynchronized activity which suggests that there typically is a “noise” component *in-vivo* that makes the spiking non-repetitive.

Although we have focused on the sensory-motor cortex of the rat and mouse, we have shown that the reconstruction framework encompasses a large spectrum from repetitive to random firing in both simulations and *in-vivo* data. Therefore, we are confident that the reconstruction framework can be applied to other data sets. Nevertheless, further verifications can be done using both whole cell patch recordings and imaging techniques. Whole cell patch recordings offer the unique advantage of providing a direct measure of the membrane potential (Neher & Sakmann 1976) and it is possible to record the simultaneous subthreshold activity in multiple neurons in the anesthetized animal (Zhao et al. 2016, Kodandaramaiah et al. 2012, Yu & Ferster 2010, Gentet et al. 2010, Volgushev et al. 2006, Lampl et al. 1999) and the awake animal (Zhao et al. 2016, Gentet et al. 2010). Imaging approaches using genetically encoded voltage indicators can record up to 4 to eight neurons simultaneously in-vivo (Adam *et al.* 2018, Piatkevich *et al.* 2019). Without simultaneous intracellular recordings it is, however, difficult to say how much of the recorded fluorescence fluctuations origin from background-, or out of focus, fluorescence from other active neurons in-vivo (Wang et al. 2018, Helmchen & Denk 2005). Finally, although imaging below 300-500 micrometer requires invasive insertion of a prism, or aspiration of brain tissue, this is irrelevant for the verification since the damaged circuitry is the same for the imaging and the reconstruction. Therefore, the reconstruction algorithm has the potential to leverage those experimental methods to address the simultaneous subthreshold dynamics in hundreds of neurons, in multiple areas, in behaving and freely moving animals with the minimal invasiveness offered by the state of the art extracellular recordings (Jun et al. 2017, Chung et al. 2019).

Finally, the algorithm was verified by comparing the reconstructed dynamics from the awake and the anesthetized state. According to previous studies the reconstructed activity is strongly correlated across layers and within layers during endogenous slow fluctuations in the anesthetized animal (Yu & Ferster 2010, Volgushev et al. 2006). In contrast, in the awake animal, we see an average correlation that is three times smaller than that of the anesthetized animal. Unfortunately, since there is no study to the authors knowledge that quantifies correlations between paired intracellular recordings in the motor cortex of the awake animal the best verification that we can find is from the overlapping sensory cortex (Harrison *et al.* 2012). Accordingly, during paw movements and during sensory stimulation, the subthreshold activity in the sensory cortex in the awake animal shows low correlations (Yu & Ferster 2010, Gentet et al. 2010, Poulet & Petersen 2008, Zhao et al. 2016). Moreover, the awake and anesthetized state differed in terms of the subthreshold dynamics. The onset of the slow fluctuations is earliest in the infragranular layers and progressively delays towards supragranular layers. Only towards the pia there seems to be an earlier onset. This pattern is common to the latency patterns shown three earlier studies. First, in the auditory cortex there is a strict monotonic increase in latencies from infragranular to supragranular layers (Sakata & Harris 2009). In a second study it is shown how the current source density has an additional earlier sink towards the pia (Manita *et al.* 2015). In the third study the onsets of slow fluctuations in intracellular recordings during anesthesia can be described by an early onset in deep layers, followed by an onset in superficial layers, and finally followed by a late onset in middle layers (Chauvette *et al.* 2010). The origin of such a progression might be the top-down projections that end in layer I and VI (Felleman & Van Essen 1991), as has been proposed by multiple authors (Manita et al. 2015, Sakata & Harris 2009). Indeed, optogenetic inhibition of axons from M2 to S1 decreases the component in S1 with the layer progression suggesting that it is manifested by a top down mechanism (Manita et al. 2015). In contrast, in the awake animal, we see the earliest onset in supragranular layers followed by the infragranular layers. This pattern resembles that in the hindlimb sensory-motor cortex (Manita et al. 2015) and only partly that of the auditory cortex (Sakata & Harris 2009). Intriguingly, some 100 milliseconds later towards a suprathreshold activation the latency difference across the layers diminishes (**Figure S3B**). This latency “cancelation” could indicate a late influence of feedback signals with the opposite latency pattern. Such a late synchronization pattern can be seen for the membrane potential for layer II/III and V neurons in the awake mouse (Zhao et al. 2016), their Fig 4D.

One millisecond resolution reconstruction was used for the laminar latency estimations. Such a resolution is only possible with a larger number of neurons since there will be more input spikes per reconstructed millisecond. The high temporal resolution might be especially beneficial for understanding the fast activity changes during for example sensory stimulation or experimentally induced perturbations. When a perturbation hits the brain the resulting activity can take many paths through the extensive neural network. Like throwing a stone in a small water pond, the water will be reflected from the uneven sides of the pond and after a very short time there will be a complex interference pattern. Given known connectivity and connection latencies a perturbation can spread across the entire brain already after 100 ms (Eriksson 2016a). This potential explosion of indirect mechanisms gives little time to record a clean effect of the perturbation. Only during the first 5 ms of a brain perturbation the effect remains in the local microcircuit. To this end it is intractable to quantify the response using spiking activity since the average firing rate of the neuron might be in the order of 1 Hz. Therefore, an instantaneous readout such as the membrane potential will enable sampling the response before it has spread outside the recorded area. Overall, large scale recordings will most likely be instrumental for studying the fast interactions that governs genesis of new brain activity in the healthy and in the diseased brain.

## Acknowledgements

Sten Eriksson for the kind contribution with a workstation.

## Author Contributions

D.E. Conceived and designed the algorithm. D.E. Analyzed the data. A.M.S. Designed and performed the simultaneous in-vivo patch and neuropixel experiments. D.E., A.M.S. and S.P. Wrote the manuscript.

## Declaration of Interests

The authors declare no competing interests.

## Methods

### Leaky integrate and fire simulation

To quantify the basic properties of the reconstruction algorithm an integrate and fire model was used. This model consisted of 200 neurons each with the following point process.

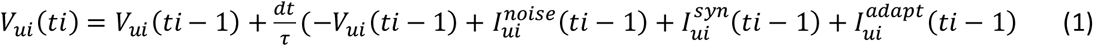

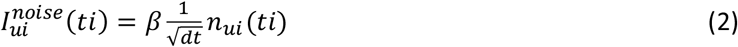

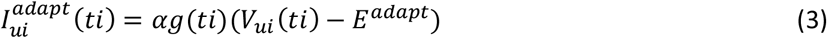

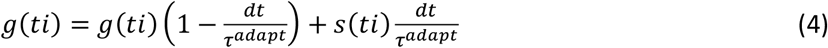

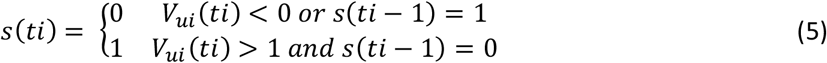

where dt=0.001, τ = 0.02, *E*^*adapt*^ = −1, *β* is the noise amplitude, and *α* = 100. The resting potential is 0. The synaptic current, 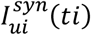, was modeled as a single dimension (i.e. combining both excitatory and inhibitory contributions and thus excluding effects of synaptic driving forces). This enabled a straightforward comparison between the reconstruction and the synaptic current, which in turn facilitated the comparison between membrane potential and synaptic input. The synaptic current was modeled as a combination of a fixed number of independent processes (If the number of independent processes is small relative to the number of recorded neurons the reconstruction becomes more defined.)

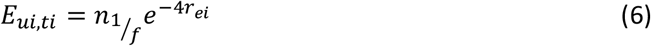

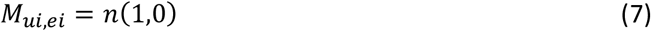

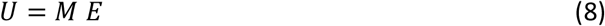

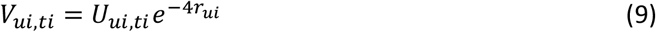

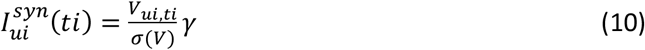

where 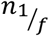 is a random distribution across time with amplitude distribution according to in-vivo membrane potential measurements in layer 2/3 (Zhao et al. 2016) which is extrapolated with a 1/f distribution for frequencies higher than 30Hz, *r*_*ei*_ (and *r*_*ui*_) is random value for each element *ei* (and *ui*) sampled from a rectangular distribution between 0 and 1, n(1,0) is a normal distribution with zero mean and unit standard deviation, *σ*(*V*) is the standard deviation across the elements in the matrix *V*, and *γ* is the connection strength. The exponents in formulas (6) and (9) result in fewer units having a large firing rate and more units having a small firing rate. The average firing rate across neurons was set to be 3 Hz. To achieve this for a certain noise amplitude, *β*, the connection strength, *γ*, had to be modified. To this end varied *β* and *γ* to achieve the following ratio, 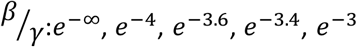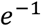, and *e*^∞^.

To test how the reconstruction depends on the repeatability of the spike trains we generated conditions for which all the trials within a given condition had an identical synaptic input (but noise was different between each trial). We tested 1, 4, 16, 256 conditions. Each trial was 4 seconds long and the trials were appended back to back.

### Processing of in-vivo data

For the awake-mouse data from Nick Steinmetz in the lab of Kenneth Harris and Matteo Carandini in the university college of London spikes were sorted automatically by Kilosort (Pachitariu et al. 2016) and manually by N. Steinmetz using Phy (Rossant *et al.* 2016). We choose to analyze data from the sensory-motor cortex. The activity in the sensory-motor cortex is typically more desynchronized than in other primary sensory areas, which on the one hand makes the reconstruction more difficult, but on the other hand gives more reliable correlation estimates (see μ-rate below).

For the simultaneous neuropixels- and patch data from the Kampff lab in the Sainsbury Wellcome Centre in London we spike sorted the neuropixels data using kilosort (Pachitariu et al. 2016). The intracellular recordings were done in sensory-motor cortices in Lister-Hooded rats of both sexes, aged between 6 weeks and 8 months, anaesthetized with urethane. The intracellular recordings where re-sampled to 30 kHz, i(t), and aligned to the neuropixels recordings (sampling rate 30kHz). To remove the spikes from the intracellular signal the spikes were detected after high pass filtering the patch data:

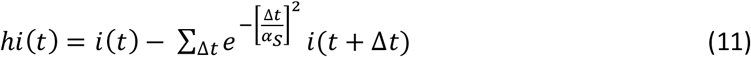

where α_S_ is 300 (10 milliseconds). The periods that satisfied *abs*(*hi*(*t*)) < 6*σ*(*hi*(*t*)) were regarded as spike-free and were assigned to a temporal mask, m(t). To increase the time around each spike this temporal mask was eroded 10 samples in time in both directions. The spikes were then replaced by 0 in the patch signal for those time points that m(t) was 0. Finally, to facilitate comparison with the reconstruction the intracellular data, i(t), was low pass filtered using the temporal mask:

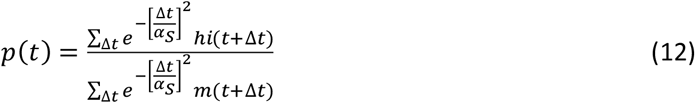

the normalization is to get an unbiased interpolation at the time points where the spikes were, m(t)=0.

### Inclusion criteria for intracellular recordings

Intracellularly recorded cells were included in the analysis if they had a signal to noise ratio larger than 10. The signal-to-noise ratio was calculated as [max(*p*(*t*)) − min(*p*(*t*))]/*σ*(p(t) − i(t)). The standard deviation, *σ*, was calculated for all time points excluding those at spikes (see methods above “Processing of in-vivo data”). It should be noted that results presented here did not qualitatively depend on the signal-to-noise threshold. The largest change is that for a smaller threshold (5) the μ-rate correlations become more similar to those of the leaky integrate and fire simulations (data not shown).

This signal to noise ratio criterion resulted in 7 cells. One cell-attached current clamped, four whole cell, and two cell attached voltage clamped. Although cell attached recordings has been used to access the membrane potential and currents in in-vitro and in-vivo (Alberio *et al.* 2018, Gonzalez *et al.* 2016, Kirmse *et al.* 2015, Perkins 2006), the disadvantage of cell attached recordings is that a small R_seal_ leads to a slower time constant (Perkins 2006). This was not an issue here since the rate of change of the fluctuations of the cell attached recordings were similar to those of the reconstructions.

### Reconstruction with neuronal network

We have used a backpropagation network for reconstructing the subthreshold activity (Figure Network). The network and its utilities were programmed in the freeware Octave. The network output is supervised by the spikes of the unit that should be reconstructed, i.e. target unit, and the network input is the spikes of all other units, i.e. source units. The feedforward propagation from input units to output/target unit does not need the target spikes, which in turns means that the reconstruction, ρ(t), is done without the knowledge of the target spikes. The feedback/backpropagation sweep needs the target spikes for training the network and optimize the weights and biases.

The spike times are binned separately for each of the U units with 10ms or 1ms time bins. The spiking data (node 16) of the target unit that we want to reconstruct is assigned to the backpropagation node (node 17) and the source spiking data is assigned to node 1 (**Figure 7**). There is a dropout after node 1 with 0.1 dropout probability. To produce a continuous signal each spike is convolved using a gated unit with one sigmoidal time constant (adjusted via backprop) in each temporal direction (forward, nodes 7-10, and backward, nodes 3-6, in time). The input units are repeated 2 times (two independent time constants) which means that there are 4 times the number of units dimensions in node 1 and throughout the network to node 11. This enables each temporal direction to be modelled by two exponentials and therefore it enables the formation of delays. Delayed filters have successfully been used to predict spike trains (Truccolo et al. 2010). The output of node 11 has 4 dimensions and contains the weighted sum (adjusted via backprop) separately for each of the four node duplicates. Each of the 4-dimensional output of node 11 is added to a bias (adjusted via backprop) and the resulting four dimensions are individually passed to a hyperbolic function (tanh). Finally, the reconstruction, ρ(t) is the weighted sum (adjusted via backprop) of the four inputs from node 14.

**Figure 7.**
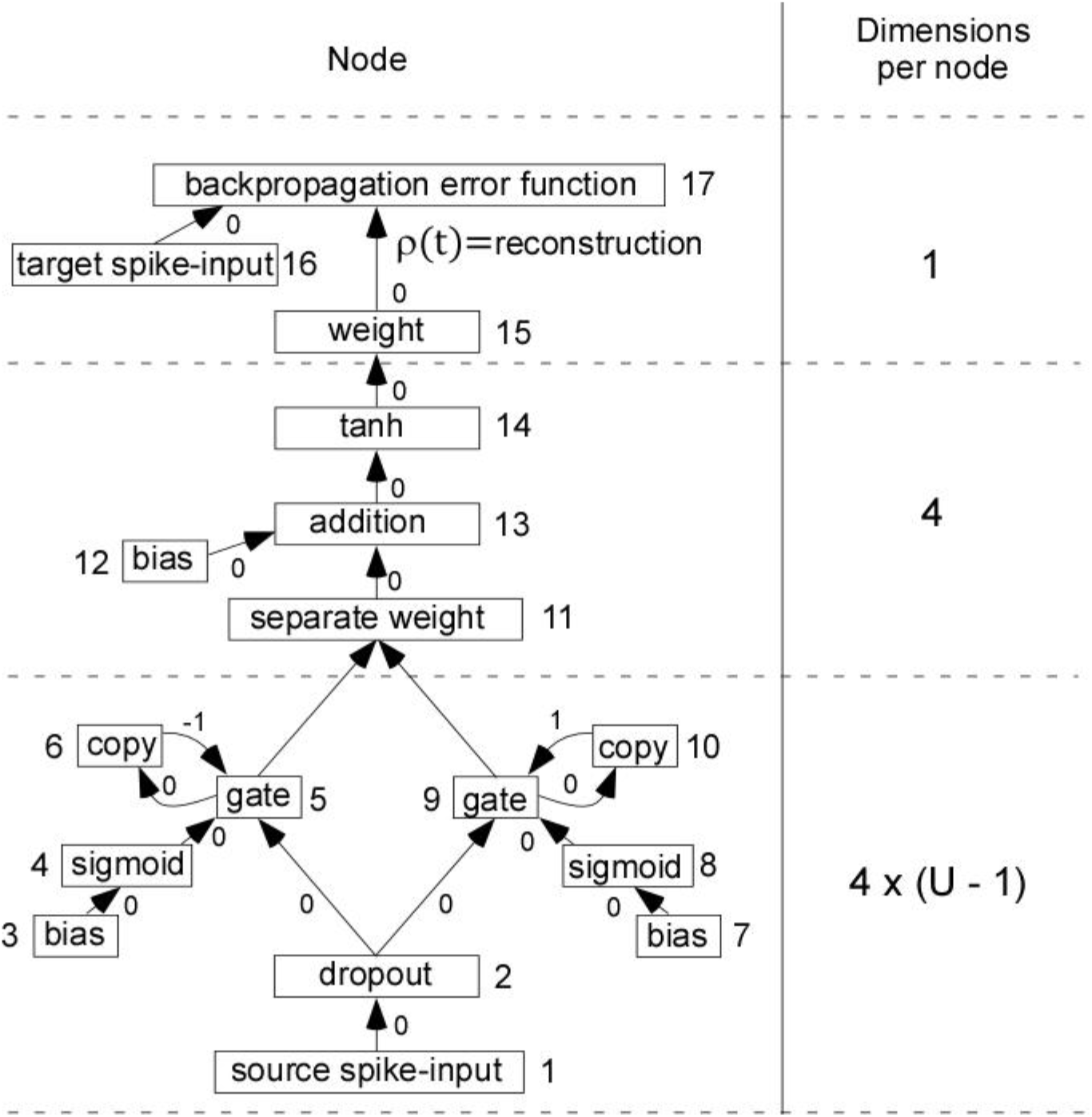
The architecture of the artificial neuronal network used to reconstruct the subthreshold activity, ρ(t). Each node corresponds to a mathematical operation (node number to the right or to the left of each node). Each arrow corresponds to the transfer of data from one node to another node, within the time sample (*0*), forward one sample (*1*) or backward one sample (*−1*).

### Backpropagation error function

A heuristic was developed for a spike-time based reconstruction error. The heuristic describes how much one gain/loose in spike time precision by moving the reconstructed activity below or above threshold. In short, if the reconstruction comes above the threshold far away from a spike then this causes a large positive error. On the other hand, if the reconstruction is below the threshold where there is a spike this causes a negative error. The spike timing balances periods of slow firing rate with periods of high firing rates without having to change or assume a specific temporal smoothing.

The error calculation is as follows. The threshold is defined at 0. First a spike probability, p(t), is created for each time bin by distributing the total number of spikes (S) to those reconstruction values, ρ(t), in node 15 that are above 0.

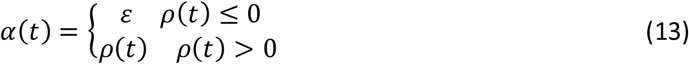

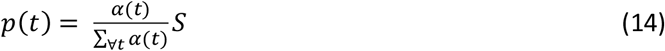

where S is the total number of spikes for the unit and *ε* is the smallest nonnegative value allowed by data type. Then the spike times are used to divide the time into regions proximal to each spike. For each spike such a region is defined by 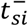 to 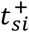.

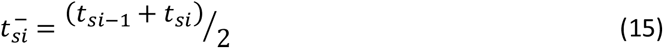

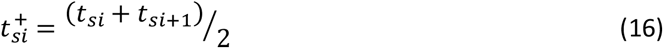

where *t*_*si*_ is the spike time of spike si. In principle the backpropagated error at a given time bin is the temporal difference between the goal spike and reconstructed spike at that time bin. But since the p(t) describes a spike probability density rather than exact spike times, the error will have the form of p(t)abs(t-*t*_*si*_). That is, at every time-bin (t) around a spike (*t*_*si*_) there will be an error p(t)abs(t-*t*_*si*_). Note that the error at the spike is p(*t*_*si*_)abs(*t*_*si*_ − *t*_*si*_) = 0. If the spike density is high long before or long after the spike, the error will be large. This positive error must be compensated by a negative error. That is, the error at the time bin of the spike must encourage the increase of the reconstruction function at that time bin (in order to cross the threshold). Afterall we want the reconstruction to be large at the time bins where there are spikes. The gain of increasing the reconstruction function at the time of a spike is the decrease in error when shifting the spike (p(t)) from a time bin (t) to that of the spike (*t*_*si*_). Therefore, the negative error at the bin of a spike is the sum of errors across all other time bins within the spike interval.

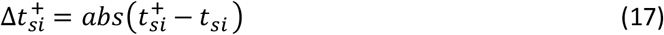

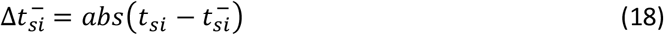

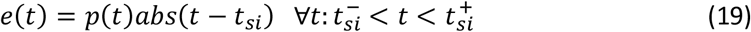

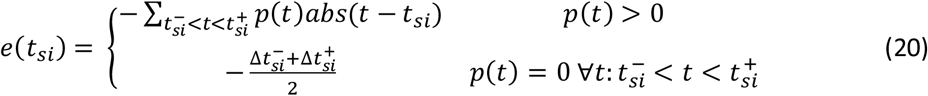

If p(t) is not above 0 at any time in the spike interval the error is the negative average of 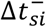 and 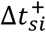. This scenario is typically the case initially before the reconstruction converges. Since the error is proportional to the inter-spike interval the error density will be constant. This in turn will make a single spike as important as a burst. Hence the algorithm will not only try to fit high firing frequencies but also single spikes and their timing are accounted for.

### Training the artificial neuronal network

Network weights (node 15 in **Figure 7**), separate weights (node 11 in **Figure 7**), were initialized with rectangular distribution 0.1+−0.01, 0+−0.1, respectively. Network gating biases (node 3 and 7 in **Figure 7**), and bias in node 12 were initialized with normal distribution 2+−0.1, and −0.1+−0.01, respectively. The network was trained with Adam with a learning rate of 0.001 (Kingma & Ba 2014).

### Comparing average instantaneous population firing rate with the reconstruction

In order to do a fair comparison between the instantaneous population firing rate and the reconstruction algorithm we smoothed the binned firing rate, r(t), with the following kernel:

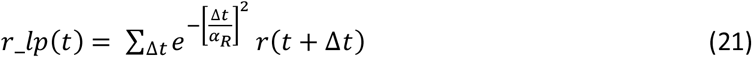

where α_R_ is 3.

### Performance indices and correlation estimation

A correlation index between the spiking data and the reconstruction was used to estimate the correlation between the reconstruction and the membrane potential. First the spike times were used to divide the time in intervals 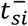 to 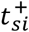 for each spike *si* (see formula above). For this interval the mean, minimum, maximum and value at the spike bin was calculated from the reconstruction.

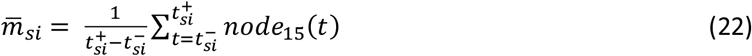

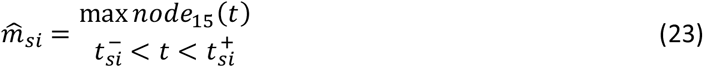

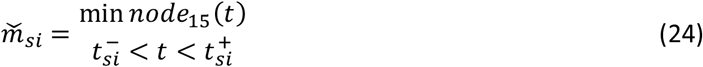

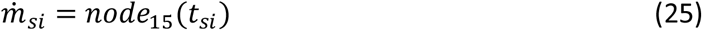

A heuristic for the correlation is then:

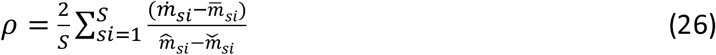

For the optimal reconstruction 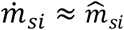, and assuming symmetric reconstruction 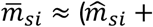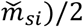, gives:

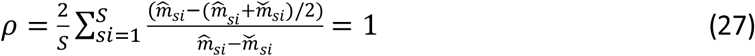

### Spike repetitiveness and μ-rate for a unit

A prerequisite for the reconstruction is that there always is at least one neuron whose firing can give information about the subthreshold activity of another neuron. The more neurons that spike while the unit, ui, is silent the better the reconstruction will be for unit ui. The μ-rate gives the instantaneous summed firing rate across all neurons at a lag relative to the unit that minimizes this summed firing rate. The rationale for the minimum operation is that any minimal instantaneous firing rate will constitute a bottleneck for the reconstruction.

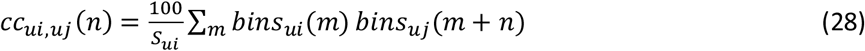

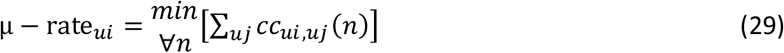

The constant 100 is there to convert the spike count to firing frequency in 10ms bins.

### Reconstruction accuracy for different number of input units

To test how the reconstruction accuracy scales with the number of input units we pseudo-randomly divided the units in groups of 2, 4, 8, 16, 29, 58, 117, 235, and 471 non-overlapping units. In each group all units were reconstructed (using the other units within that group as input units). The number of groups (and the total number of units) were 235 (470), 117 (468), 58 (464), 29 (464), 16 (464), 8 (464), 4 (468), 2 (470), and 1 (471) units. The number of input units in each group was 1, 3, 7, 15, 28, 57, 116, 234, and 470.

When comparing the averaged correlation indices across the two largest groups (the groups with 234 and 470 input units) we noticed that the average correlation did not increase from the smaller to the larger group as would be expected. This could be because the algorithm did not get enough data (spikes) to fit its parameters. Therefore, we tested if the average correlation would increase if we only reconstructed units with a high spike count:

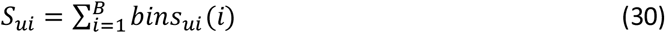

where B is the number of 10 millisecond bins in the recording.

### Calculating latencies across cortical layers

To calculate the latency across cortical layers we first calculated the time points where the average reconstructed activity across units, a(t) passed a certain threshold. For the anesthetized recordings where there were clear transitions between down and up states, we used the temporal average of a(t) to define the threshold. For the awake recordings where we did not have clear state transitions, we calculated a bandpass filtered version using a low and high pass filter according to the following formula:

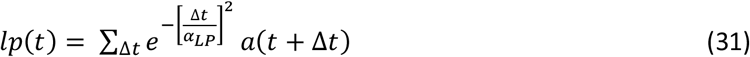

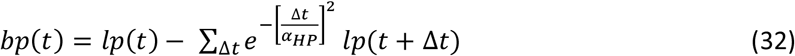

where α_*LP*_ = 100 ms and α_*HP*_ = 400 ms. The mean of the bandpass filtered signal, bp(t), served as a threshold. A certain temporal window (−100ms to 100ms) around those threshold crossings were used to create a threshold triggered average of the reconstructed activity. This resulted in a two-dimensional matrix with 471 rows (units) and 201 columns (time samples around each threshold crossing). Three latencies for each unit was extracted by taking the time sample of the 10%, 50%, and 100% of the maximum value.

The variability of the latency was estimated using bootstrapping. To this end threshold crossings were sampled with replacement to conserve the number of threshold crossings and the latencies was calculated for the corresponding threshold triggered average of the reconstructed activity. This procedure was repeated 100 times and the standard deviation was calculated for the resulting latencies (**Figure 6C**). To minimize the influence of outliers in this estimate we also smoothed the reconstructed activity across neighbouring units (**Figure 6D**). The filter used was:

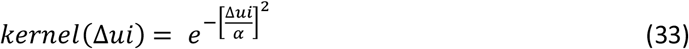

where α is 35.

### Calculating the dimensionality of the activity in each layer

The mean reconstructed activity was subtracted from respective unit. Then a singular value decomposition (SVD) was calculated from the sum-of-squares-and-cross-products matrix (summed across time points) for the normalized reconstructed activity for the units in each layer. The dimensionality for the layer was then taken as the number of eigenvalues that are larger than 10% of the largest eigenvalue. To get the variability of this dimensionality we applied bootstrapping. The SVD was applied for a new set of units taken with replacement in order to conserve unit count. This procedure was repeated 100 times and the standard deviation was calculated for the resulting dimensionalities.

## Notes

https://github.com/David-Eriksson/SubLab

