## Supplementary material for "Reconstruction of in-vivo subthreshold activity of single neurons from large-scale spiking recordings": Figure S1

[*]) Integrative Medical Biology (IMB)
Umeå University
90187 Umea
Sweden

[**]) Sainsbury-Wellcome Centre for Neural Circuits and Behaviour
25 Howland Street
London W1T 4JG
England

[***]) Zentrum für Neurowissenschaften (ZfN)
Albertstr. 23
79104 Freiburg
Germany


[+]) corresponding author

**Supplemental Figures**

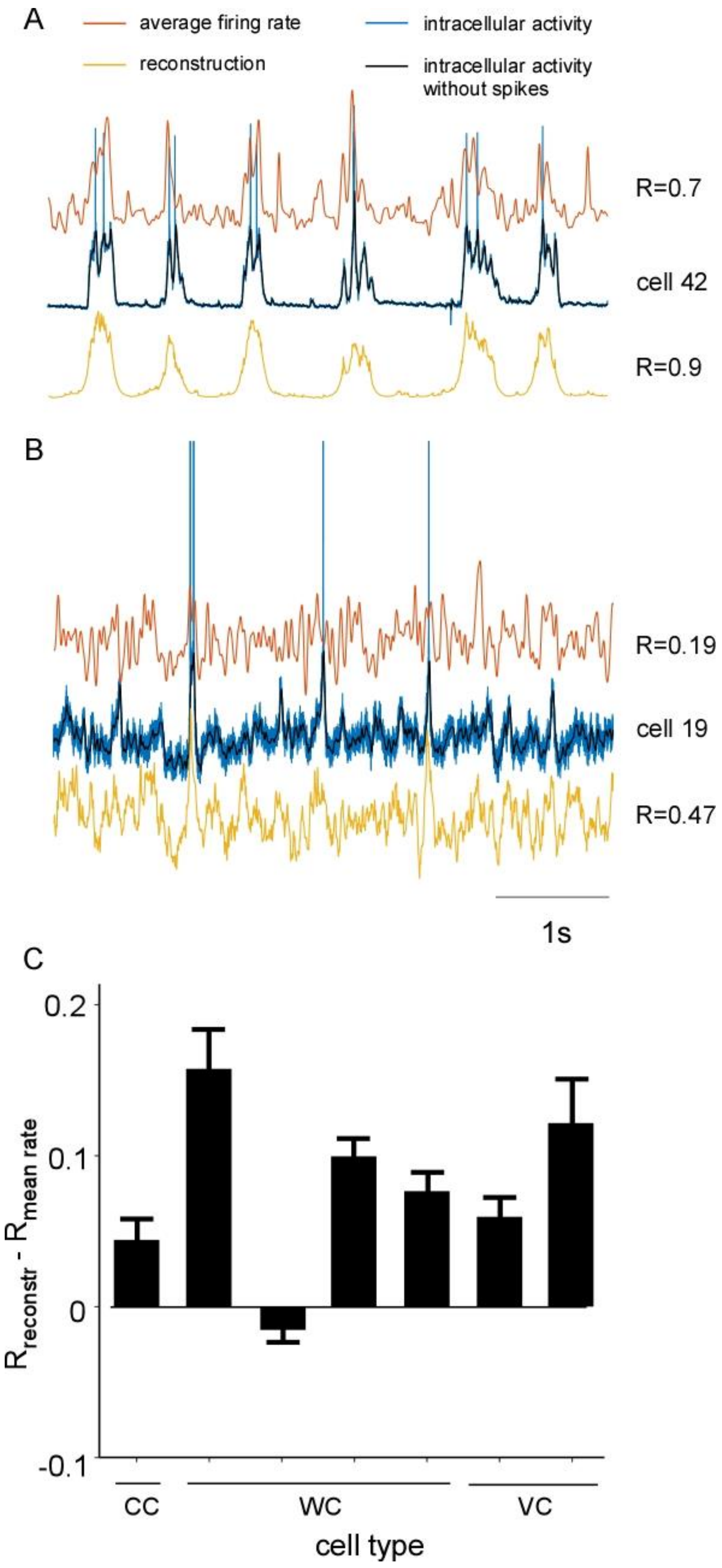


**Figure S1**. The reconstruction algorithm outperforms, in terms of correlation with the true recordings, the average instantaneous firing rate across units, related to **Figure 2**. A: Reconstruction (yellow) of a trace with pronounced UP/DOWN states (blue) and the average instantaneous firing rate across neurons (red). **B**: Reconstruction of a more desynchronized trace and the average instantaneous firing rate across neurons. **C**: Statistical comparison between the correlation of the reconstruction and the ground truth recordings and instantaneous firing rate and ground truth recordings across all 7 cells. Each cell was reconstructed at 10 (except for the current clamped cell which only had 7 because of a shorter recording) different test intervals (as opposed to the remaining time points that served as training). This gave 10 (or 7) values for each cell for which the standard deviation of the mean was calculated.
