## Supplementary material for "Reconstruction of in-vivo subthreshold activity of single neurons from large-scale spiking recordings": Figure S2

[*]) Integrative Medical Biology (IMB)
Umeå University
90187 Umea
Sweden

[**]) Sainsbury-Wellcome Centre for Neural Circuits and Behaviour
25 Howland Street
London W1T 4JG
England

[***]) Zentrum für Neurowissenschaften (ZfN)
Albertstr. 23
79104 Freiburg
Germany


[+]) corresponding author

**Supplemental Figures**

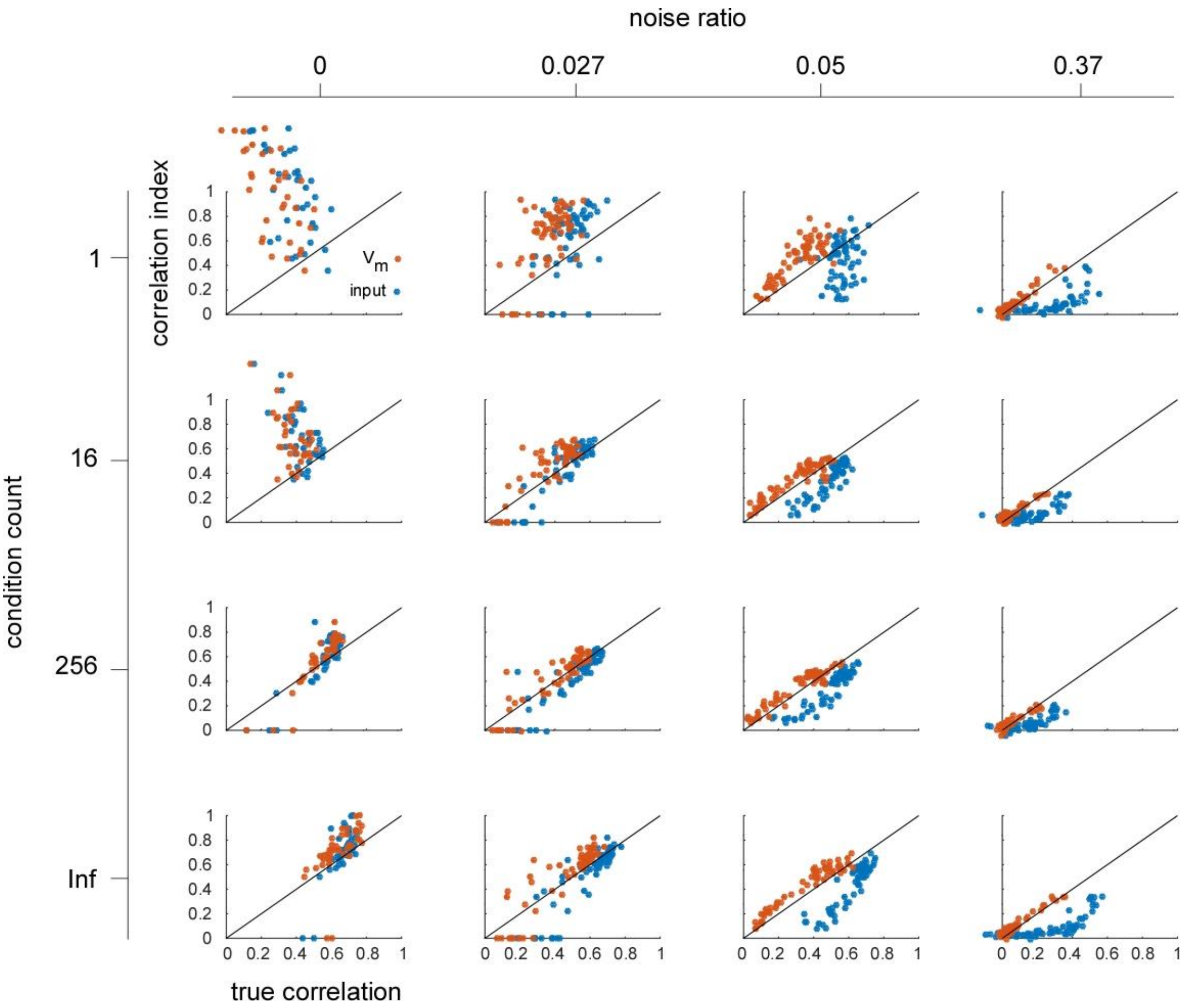


**Figure S2**. Relation between true correlation and correlation index for different common input strengths and noise ratios, related to **Figure 3**. The correlation index underestimates the correlation with the synaptic input when the noise ratio is large. Since the noise-jitter is independent across the neurons the reconstruction algorithm can cancel-out this jitter thereby giving a large true correlation between the reconstruction and the synaptic input. In contrast, the noise jitters the spikes which in turn causes a low correlation between spikes and reconstruction (correlation index). Therefore, if we see an underestimation of the true correlation this can be because there is a spike jitter that is independent across the neurons and independent of the reconstructed signal.
