## Supplementary material for "Reconstruction of in-vivo subthreshold activity of single neurons from large-scale spiking recordings": Figure S3

[*]) Integrative Medical Biology (IMB)
Umeå University
90187 Umea
Sweden

[**]) Sainsbury-Wellcome Centre for Neural Circuits and Behaviour
25 Howland Street
London W1T 4JG
England

[***]) Zentrum für Neurowissenschaften (ZfN)
Albertstr. 23
79104 Freiburg
Germany


[+]) corresponding author

**Supplemental Figures**

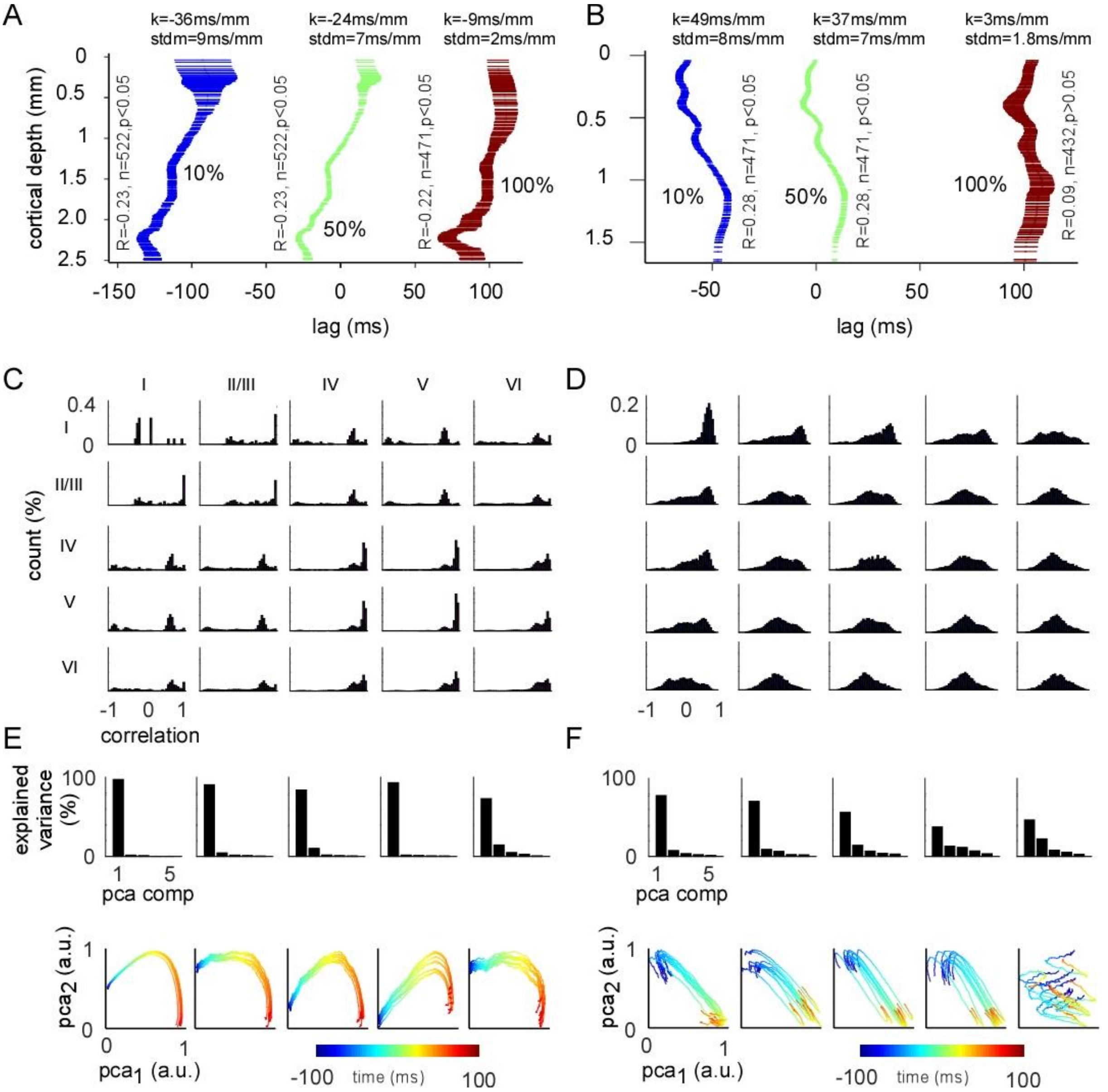


**Figure S3.** Comparison of reconstructed activity for different activity levels (**A**-**B**) and for different layers (**C**-**F**) in the anesthetized and awake animal, related to **Figure 6**. **A:** Based on the rise triggered average of reconstructed activity for anesthetized recordings (see **Figure 6C)**. The activity was smoothed across neighboring units (see methods), and then latency at 10% (blue), 50% (green), and 100% (red) of the maxima was calculated for each unit (right). Bars indicate the standard deviation of the bootstrapped latencies. Note that the correlation statistics is for the latency as a function of cortical depth (not vice versa). **B:** Same as in **A** but for awake recordings. **C:** Distribution of cross-correlation values for unit-pairs for each combination of layers. The location of the cortical layers has been adopted from Yamawaki et. al. (Yamawaki *et al.* 2014). On the diagonal are the distribution of the cross-correlation for all unit pairs within a certain layer. **D:** Same as in **C** but for awake recordings. **E:** The principal component analysis for the units for a certain layer for the anesthetized recordings. Only the five first components are shown for clarity (top). Ten bootstrapped trajectories (sampled with replacement across the threshold crossings) of threshold crossing triggered reconstructed activity (see **Figure 6C**) along the first two principal components. Time along the trajectory is coded with color from blue to red (bottom). **F:** Same as in **E** but for awake recordings.
